# Siderophore from *Talaromyces trachyspermus:* augmentation and characterization

**DOI:** 10.1101/2021.04.13.439607

**Authors:** Sharda Sahu, Anil Prakash

## Abstract

In the present study, a siderophore compound produced by an endophytic fungus, *Talaromyces trachyspermus* was optimized for maximum production, 88.9 % SU by applying Plackett-Burman design and Response Surface Methodology through Central Composite Design that showed the succinic acid (1.141 g/L), sucrose (31.028 g/L) and temperature (27.475 °C) as significant factors. On scale up, a further increase in siderophore yield was obtained (by 3%) The compound was extracted, purified and detected chemically as catecholate siderophore showing max. λ absorbance at 279nm. Contained of hydroxy benzene as shown by GC-MS analysis and further identified as berberine by HRLC-MS studies. The compound berberine is clinically a very important drug with several ethnobotanical properties. This is rare to report fungal catecholate and first to report the production of berberine from *Talaromyces species*.

## Introduction

Most filamentous ascomycete fungi biosynthesize high affinity iron chelators called siderophores, a low molecular weight secondary metabolite, produced by microbial non ribosomal peptide synthetases (NRPSs) under iron deficit, to stream iron to the organism. Iron, which has readily inter-convertible oxidation states, Fe (II) and Fe (III) is the fourth most abundant earth element and is greatly needed by most microorganisms in order to perform several metabolic functions such as redox reactions, heme synthesis and electron transport during ATP synthesis [1, 2]. It acts as cofactor for several enzymes [3]. Naturally occurring iron compounds are insoluble in water and this small amount of available iron in nature acts as growth-limiting factor for several living organisms [4].

Microbial siderophores may be grouped as hydroxamates, catecholates, carboxylates, or mixed type depending upon the chemical nature of their coordination sites [5]. Being iron-chelators, they have established much attention in agricultural, environmental, medicinal and biotechnological applications. They also perform key functions in fungal cells by aiding in sexual and asexual spore germination and development by storing iron reserves within spores as studies revealed in fungus such as *Neurospora crassa, Penicillium chrysogenum, Aspergillus nidulans, Cochliobolus heterostrophus* and *Fusarium graminearum* [6]. Intracellular siderophores are understood to buffer against reactive oxygen species (ROS) generated by the Haber-Weiss-Fenton reaction in the presence of unbound iron, by seizure of cellular free iron. Under laboratory conditions, these siderophores can be produced and optimized in liquid medium through statistical augmentation as the process and pave an ideal way for optimization of several biochemical and biotechnological processes [7]. Ensuing, they can be detected and identified by different methods.

An attempt has been made for production and optimization of physiochemical parameters of *Talaromyces trachyspermus*, an endophyte, isolated from foliar tissues of *Withania somnifera* and exhibited comparatively higher activity for hydrolytic enzymes, protease, chitinase, amylase, cellulase and pectinase that are required for antagonistic property. It was observed to be a propitious and proxy biocontrol agent against plant pathogen, *Sclerotinia sclerotiorum*. This strain is also characterized with IAA (69 ± 0.05), siderophore (80%SU) and phosphate solubilization (550 ± 0.4 µg/ml) and showed inhibition of phytopathogen (19 mm zone of inhibition in 24 h) [8]. Present work focuses on statistical optimization approaches, i.e., Plackett–Burman (PB) design and response surface methodology (RSM) by central composite design (CCD), for enhanced production of siderophore by using Czapek’s Dox broth medium supplemented with leucine, biotin, and succinic acid. Scale-up of optimized shake flask protocol to laboratory-scale bioreactor and characterization of compound was also studied.

## Materials and Methods

### Optimizationofphysicalandchemicalparametersforsiderophoreproduction

#### Statistical approach & Designs of experiment

In the first phase, Plackett-Burman design was used to find the significant variable(s) or factor(s) to optimize the siderophore production. In the next phase, Response Surface Methodology through Central Composite Design was used to find the optimum concentration of the selected variable. According to Murugappan et al. the selected variables on the responses was analyzed to maximize the siderophore production [9].

#### Plackett–Burman design

At first stage of the optimization, 13 variable Plackett-Burman experimental design was used to find the significant ingredients of the basal medium, Czapek’s Dox broth for the maximum production of siderophore. This experimental design was two-factorial design, and was used to identify the critical parameters required for optimum siderophore production by screening n variables in n + 1 experiment [10]. Optimum values for physical parameters such as pH, temperature, RPM and incubation days were obtained by screening one variable at a time method (OVAT) (Table 1). These variables were coded as A, B, C, D, E, F, H, J, K, L, M, and N respectively, while O, P, Q, R, S and T were considered as dummy variables. Experiments of PB design were carried out in triplicates, and the average mean of siderophore unit percentage (SU%) was considered as the final response. The quantification of siderophore production was carried out by Chrome Azurole Shuttle assay [11]. Each of these variables was represented at two different levels-high concentration (H) and low concentration (L) (Table 1). The effect of individual variable was determined by calculating the difference between the average of measure at the H value and the L value. The full experimental design consisting of 13 variables with six dummy variables and 20 trials is depicted in table 2. The effect of each variable on siderophore production in the form of SU% was calculated, and the highest confidence level and F-test was used to determine the significant component [12].

**Table 1:**
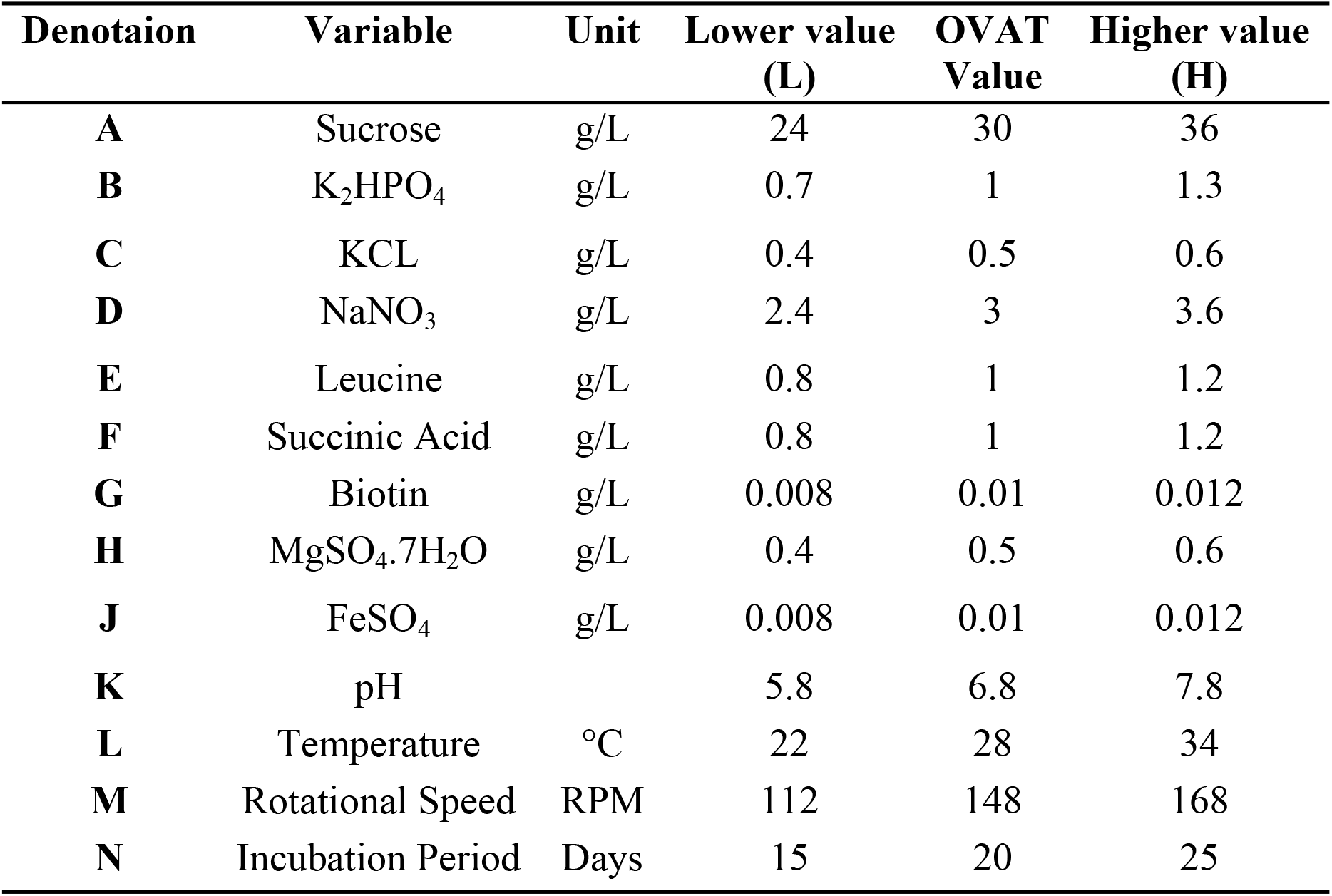
Variables and their values in Plackett-Burman Design for siderophore production by *T. trachyspermus*.

| Denotaion | Variable | Unit | Lower value<br>(L) | OVAT<br>Value | Higher value<br>(H) |
| --- | --- | --- | --- | --- | --- |
| **A** | Sucrose | g/L | 24 | 30 | 36 |
| **B** | K<sub>2</sub>HPO<sub>4</sub> | g/L | 0.7 | 1 | 1.3 |
| **C** | KCL | g/L | 0.4 | 0.5 | 0.6 |
| **D** | NaNO<sub>3</sub> | g/L | 2.4 | 3 | 3.6 |
| **E** | Leucine | g/L | 0.8 | 1 | 1.2 |
| **F** | Succinic Acid | g/L | 0.8 | 1 | 1.2 |
| **G** | Biotin | g/L | 0.008 | 0.01 | 0.012 |
| **H** | MgSO<sub>4</sub>.7H<sub>2</sub>O | g/L | 0.4 | 0.5 | 0.6 |
| **J** | FeSO<sub>4</sub> | g/L | 0.008 | 0.01 | 0.012 |
| **K** | pH |  | 5.8 | 6.8 | 7.8 |
| **L** | Temperature | °C | 22 | 28 | 34 |
| **M** | Rotational Speed | RPM | 112 | 148 | 168 |
| **N** | Incubation Period | Days | 15 | 20 | 25 |

**Table 2:**
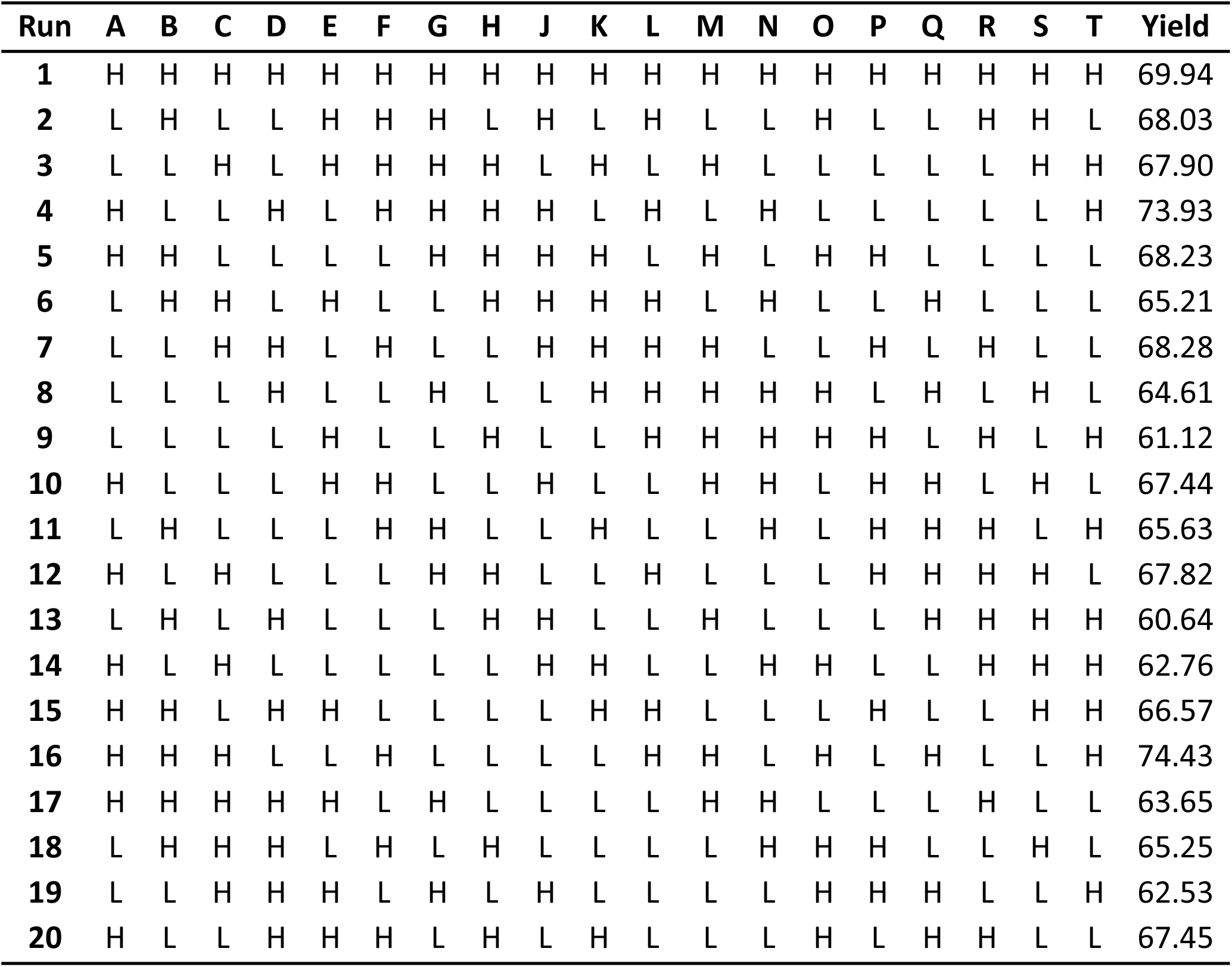
Variables placket-Burman experimental design for siderophore production by *T. trachyspermus*.

#### Response surface methodology (RSM)

The second phase in optimization of the medium-composition was to find the optimum concentration of the significant components by RSM through CCD. It involved steps such as procedures to find the optimum region, the responses in the optimum region of variables, estimation of the optimal conditions and verification of the data. The variables obtained from PB design to enhance the siderophore production were selected for CCD to study the interaction between the various medium constituents, which influence the siderophore production.

In the present work, experiments were planned to obtain a quadratic model. Hence, the concentrations of the three factors i.e., sucrose, succinic acid and temperature, identified by PB design, were optimized, keeping other variables constant at zero (0) level. Each factor was studied at five different levels (-**α**, −1, 0, +1, +**α**). The complete experimental plan of CCD with respect to their values in coded and actual form is listed in table 3. SU% was measured in triplicate in 20 trial-experimental runs. Design matrix consists of coded terms with eight factorial points, six axial points and five central points. The data were fitted into the second-order polynomial equation, and the coefficients were calculated and analyzed. The general form of the second-degree polynomial equation is:

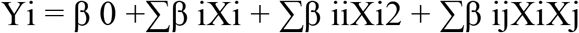

**Table 3:** Analysis of Placket-Burman design for siderophore production by *T. trachyspermus*.

| Denotation | Source | Sum of squares | Mean Square | F Value | P value |
| --- | --- | --- | --- | --- | --- |
| A | Sucrose | 54.51 | 54.51 | 1186000.0 | 0.001 |
| B | K<sub>2</sub>HPO<sub>4</sub> | 0.70 | 0.70 | 15179.21 | 0.005 |
| C | KCl | 0.85 | 0.85 | 18555.07 | 0.005 |
| D | NaNO<sub>3</sub> | 1.64 | 1.64 | 35679.73 | 0.003 |
| E | Leucine | 6.90 | 6.90 | 150000.00 | 0.002 |
| F | Succinic Acid | 101.82 | 101.82 | 2214000.0 | 0.000 |
| G | Biotin | 8.64 | 8.64 | 188000.00 | 0.002 |
| H | MgSO<sub>4</sub>.7H<sub>2</sub>O | 0.63 | 0.63 | 13739.48 | 0.005 |
| J | FeSO<sub>4</sub> | 0.32 | 0.32 | 6968.85 | 0.008 |
| K | pH | 0.15 | 0.15 | 3319.97 | 0.011 |
| L | Temperature | 40.50 | 40.50 | 880900.00 | 0.001 |
| M | Rotational Speed | 0.06 | 0.06 | 1213.72 | 0.018 |
| N | Incubation Period | 7.62 | 7.62 | 165800.00 | 0.002 |
| O | O | 0.37 | 0.37 | 7994.71 | 0.007 |
| P | P | 1.68 | 1.68 | 36486.54 | 0.003 |
| Q | Q | 0.00 | 0.00 | 0.00 | 0.999 |
| R | R | 21.60 | 21.60 | 469800.00 | 0.001 |
| S | S | 4.53 | 4.53 | 98588.96 | 0.002 |
| T | T | 0.01 | 0.01 | 290.40 | 0.037 |
|  | Model | 252.53 | 13.29 | 289079.82 | 0.001 |
|  | Residual | 0.00 | 0.00 |  |  |
|  | Cor Total | 252.53 |  |  |  |

Where Yi is the predicted response, XiXj are input variables, which influence the response variable Y, β 0 is the offset term, βi is the ith linear coefficient, β ii is the quadratic coefficient, β ij is the ijth interaction coefficient. After the analysis of data, the experimental run with optimum values of variable was done to check the validity of the model.

#### Scale-up and validation on bioreactor

The shake flask process was scaled-up to 5L capacity fully automated laboratory-scale bioreactor (Murhotpye Scientific Co., India, Model LF-5) equipped with monitoring & control of physiochemical variables, namely, pH temperature, dissolved Oxygen, air flow rate & agitation. The fermenter was operated at 3 L working volume. The fermenter was autoclaved and sterilized at 121°C for 20 minutes. Thereafter, separately autoclaved/ filter-sterilized components of medium were added to bioreactor. Initial value of the physicochemical parameters was adjusted (aeration rate of 0.1vvm, pH at 6.8, temperature 28°C, agitation 140 rpm). Following inoculation of the medium, the DO was automatically controlled with respect to aeration and agitation. The pH was adjusted by automatic addition of 2N KOH/2N H_3_PO_4_. The performance of siderophore producing isolate was tested out and the validity of optimization studies carried out at shake-flask level was confirmed.

#### Characterization of siderophore

##### Chemical detection of type of siderophore

Detection of hydroxamate type of siderophores was carried out using the Csaky Assay [13]. The assay detects the presence of secondary hydroxamate and depends on the oxidation to nitrite and formation of a colored complex via diazonium coupling. The reaction mixture after incubation of 30 minutes, a cherry color indicated the presence of hydroxamate type of siderophore. And to check the presence of catechol type siderophore, Arnow assay was performed [14].

##### Extraction and purification of siderophore

Cell free supernatant was harvested and CAS positive supernatant of the fungi was concentrated (10X) on rotatory vacuum evaporator (Model) at 100 rpm at 50° C (pH 6.0 adjusted with 12 N HCl). Then it was subjected to Amberlite XAD column prepared with AmberliteXAD-16 resin (Sigma, USA) dissolved in distilled water and kept overnight at 8°C for soaking. Before use, the resin was activated by treating with 0.1% NaNO_3_ and loaded in a sintered glass column of 40×6cm size; the loaded column was pre-washed twice with water, methanol and water.

The aqueous concentrated supernatant containing ferric siderophore was allowed to slowly pass through the column at the rate of 5ml/min. using a peristaltic pump. Loading of supernatant was continued until the saturation of column, as indicated by reddish-browning of column. The column was then washed with five to ten bed volume of distilled water to remove all unbound components of the medium. Solvent was then changed to 50% methanol and the column was left to equilibrate for the extraction of siderophores. Different fractions were separately collected; filtrate, water and eluted fractions were checked for CAS test. CAS positive fractions were used for characterization of siderophore.

##### Detection Using Thin Layer Chromatography (TLC)

CAS positive fraction or concentrated samples of siderophore were spotted on HiMedia 5×10 silica gel plates and spots were allowed to dry. The plates were then run in an n-butanol: acetic acid: DH2O (12:3:5) solvent system until the solvent front reached the top of the plate. Plates were then dried and sprayed with 0.1 M FeCl_3_ in 0.1 N HCl. The formation of a wine-colored spot indicated a hydroxamate-type siderophore, while a dark gray spot indicated production of a catechol-type siderophore. Siderophores were separated on the basis of hydrophobicity using these plates.

##### Spectrophotometric assay

CAS positive fraction was subjected to double beam UV-Visible Spectrophotometric scan (200-800 nm) and the absorption maxima were recorded.

##### Fourier Transform Infrared (FTIR) spectroscopy

The extracted siderophores were characterized by collecting the FT-IR spectra in the transmission mode (4000-450/cm) using Shimadzu FT-IR spectrophotometer (Model-8400S and IR solution version 1.40 software).

##### GC-MS analysis

The GC-MS analysis was performed at IISERB-CIF-Mass Facility Bhopal (M.P) using an Agilent GC-MS system (7890A GC with 5975C MS). Samples (1 µl volume) were injected in split-less mode. An Agilent Technologies HP-5MS (30 min by 0.250 mm by 0.25 min) fused silica capillary column was used.

##### HRLC-MS analysis

HRLC-MS/MS analysis was performed at Sophisticated Analytical Instrument Facility (SAIF) Indian Institute of Technology, Bombay using electrospray ionization mass spectrometer (ESIMS) with a quadrupole time-of-flight (Q-TOF) mass spectrometer (model G6550A). Separation of compound was done using the Luna(r) 5um C18 150 x 2mm (Phenomenex). Mobile phase A consisted of 0.1% formic acid in water while mobile phase B was 0.1% formic acid in acetonitrile. A gradient separation, using 10%–90% mobile phase B was performed at a flow rate of 0.2 ml/min. The injection volume was 6 μ L and the column temperature was set at 40°C. The eluted compound was injected directly into the tandem quadrupole mass spectrometer operated in the positive electrospray ionization (ESI+) mode HRLC-MS/MS analysis was performed HRLC-MS/MS analysis was performed a capillary voltage of 1000V with nitrogen gas at a temperature of 250°C. Data was acquired in the multiple reaction monitoring (MRM) mode and 6200 series TOF/6500 series Q-TOF B.05.01 (B5125) software was used for control of the equipment and data acquisition.

## Results and Discussion

To allow an economical use of siderophore sand metabolic resources, the need of chemical stability and regulation of fungal siderophore biosynthesis is an essential aspect of microbial life in the soil and in the rhizosphere for determination of significant variables affecting the response of siderophore production. Statistical optimization of these variables for maximum production was carried out.

Medium selected for analysis of *T. trachyspermus* was Czapek’s Dox Broth, which contains sucrose, sodium nitrate, dipotassium phosphate, magnesium sulfate, potassium chloride and Iron sulfate in the media. FeSO_4_is already present as iron source, so there was no requirement of putting the external iron source for the study. Along with these variables and external variables, a total of 13 factors were considered for Placket-Burman analysis, which have been listed in table 2 with their high and low value for Placket Burman design. The results of the PBD analysis have been depicted in table 3. The three significant variables having a major effect on siderophore production with *T. trachyspermus* were found to be succinic acid, sucrose and temperature as deduced from F value and Pareto chart (Figure 1) of the effect of a variable on siderophore production. The model F-value of 289079.82 implies that model is significant. There remains merely 0.14% chance for F-value to be large occurring due to noise.

**Fig 1.**
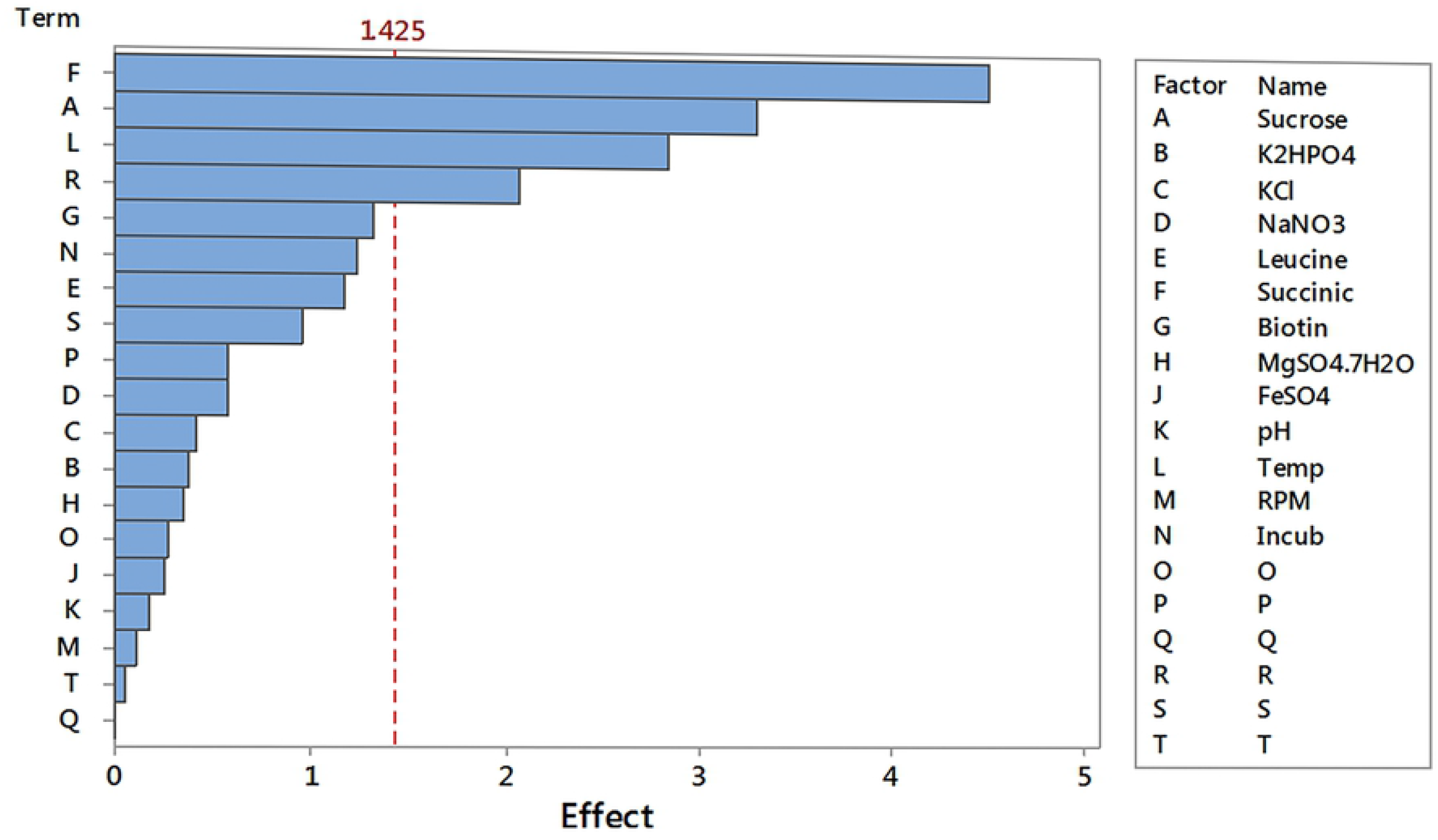
Pareto chart for Placket-Burman analysis for Siderophore production in *T. trachyspermus*.

During RSM run, the value of significant variables obtained in PBD varied as shown in table 4 while keeping all other variables at their zero-value obtained during testing one variable at a time. There were 20 runs in triplicate as shown in table 5.

**Table 4:** Variables and their values for Response Surface Methodology for siderophore production by *T. trachyspermus*.

| Denotation | Variable | Unit | -α | -1 | 0 | 1 | A |
| --- | --- | --- | --- | --- | --- | --- | --- |
| A | Succinic Acid | g/L | 0.6636 | 0.8 | 1 | 1.2 | 1.3364 |
| B | Sucrose | g/L | 19.9092 | 24 | 30 | 36 | 40.0908 |
| C | Temperature | g/L | 17.9092 | 22 | 28 | 34 | 38.0908 |

**Table 5:** Response Surface Methodology Design for siderophore production by *T. trachyspermus*.

| A | B | C | Experimental | Predicted |
| --- | --- | --- | --- | --- |
| 1 | -1 | -1 | 72.65 | 75.33 |
| 1 | 1 | 1 | 77.88 | 78.75 |
| -1 | 1 | -1 | 63.99 | 65.19 |
| $\alpha$ | 0 | 0 | 84.68 | 83.58 |
| 0 | $-\alpha$ | 0 | 60.99 | 60.76 |
| 0 | 0 | 0 | 86.24 | 86.03 |
| 0 | 1 | -1 | 75.93 | 78.21 |
| $-\alpha$ | 0 | 0 | 57.93 | 58.51 |
| -1 | -1 | -1 | 59.03 | 59.09 |
| -1 | 1 | 1 | 66.92 | 65.18 |
| 0 | 0 | 0 | 85.87 | 86.03 |
| 0 | 0 | $\alpha$ | 73.59 | 75.51 |
| -1 | -1 | 1 | 60.12 | 59.92 |
| 0 | $\alpha$ | 0 | 70.74 | 69.63 |
| 0 | 0 | 0 | 86.30 | 86.03 |
| 1 | -1 | 1 | 75.70 | 74.30 |
| 0 | 0 | 0 | 86.75 | 86.03 |
| 0 | 0 | 0 | 85.95 | 86.03 |
| 0 | 0 | $-\alpha$ | 79.65 | 76.39 |
| 0 | 0 | 0 | 85.68 | 86.03 |

The second-degree polynomial obtained for yield from RSM analysis for *T. trachyspermus* was: Yield = 86.03+7.45* A+2.63* B-0.26* C-0.20* AB-0.46* AC −0.21* BC-5.30* A^2-7.36* B^2-3.56* C^2. The Model F-value of 59.27 implies that the model is significant. There remains merely 0.01% chance for F-value to be large occurring due to noise. Values of Prob> F less than 0.0500 indicate model terms are significant. The Lack of Fit F-value of 50.91 implies the Lack of Fit is significant. There remains merely 0.03% chance that a Lack of Fit F-value to be large occurring due to noise fig 2 and fig 3 show the contour plot and surface plot respectively of the RSM analysis (Table 6), each considering one variable being at zero value.

**Table 6:** Analysis of variance by Response Surface Methodology for siderophore production by *T. trachyspermus*.

| Source | Some of Squares | df | Mean Square | F Value | Prob>F |  |
| --- | --- | --- | --- | --- | --- | --- |
| Model | 2007.430 | 9 | 223.050 | 59.270 | < 0.0001 | Significant |
| A-NH<sub>4</sub>Cl | 610.237 | 1 | 610.237 | 158.317 | 1.9E-07 |  |
| B-Glucose | 17.364 | 1 | 17.364 | 4.505 | 0.05977 |  |
| C-Succinic Acid | 13.202 | 1 | 13.202 | 3.425 | 0.09394 |  |
| AB | 0.142 | 1 | 0.142 | 0.037 | 0.85184 |  |
| AC | 4.287 | 1 | 4.287 | 1.112 | 0.31643 |  |
| BC | 0.905 | 1 | 0.905 | 0.235 | 0.63836 |  |
| A<sup>2</sup> | 348.250 | 1 | 348.250 | 90.349 | 2.5E-06 |  |
| B<sup>2</sup> | 1145.488 | 1 | 1145.488 | 297.181 | 9.1E-09 |  |
| C<sup>2</sup> | 172.967 | 1 | 172.967 | 44.874 | 5.4E-05 |  |
| Residual | 38.545 | 10 | 3.855 |  |  |  |
| Lack of Fit | 37.136 | 5 | 7.427 | 26.345 | 0.00134 | Significant |
| Pure Error | 1.410 | 5 | 0.282 |  |  |  |
| Cor Total | 2250.587 | 19 |  |  |  |  |
| Std. Dev. | 1.963 |  | R-Squared | 0.983 |  |  |
| Mean | 66.317 |  | Adj R-Squared | 0.967 |  |  |
| C.V. % | 2.960 |  | Pred R-Squared | 0.857 |  |  |
| PRESS | 321.042 |  | Adeq Precision | 19.682 |  |  |

**Fig 2.**
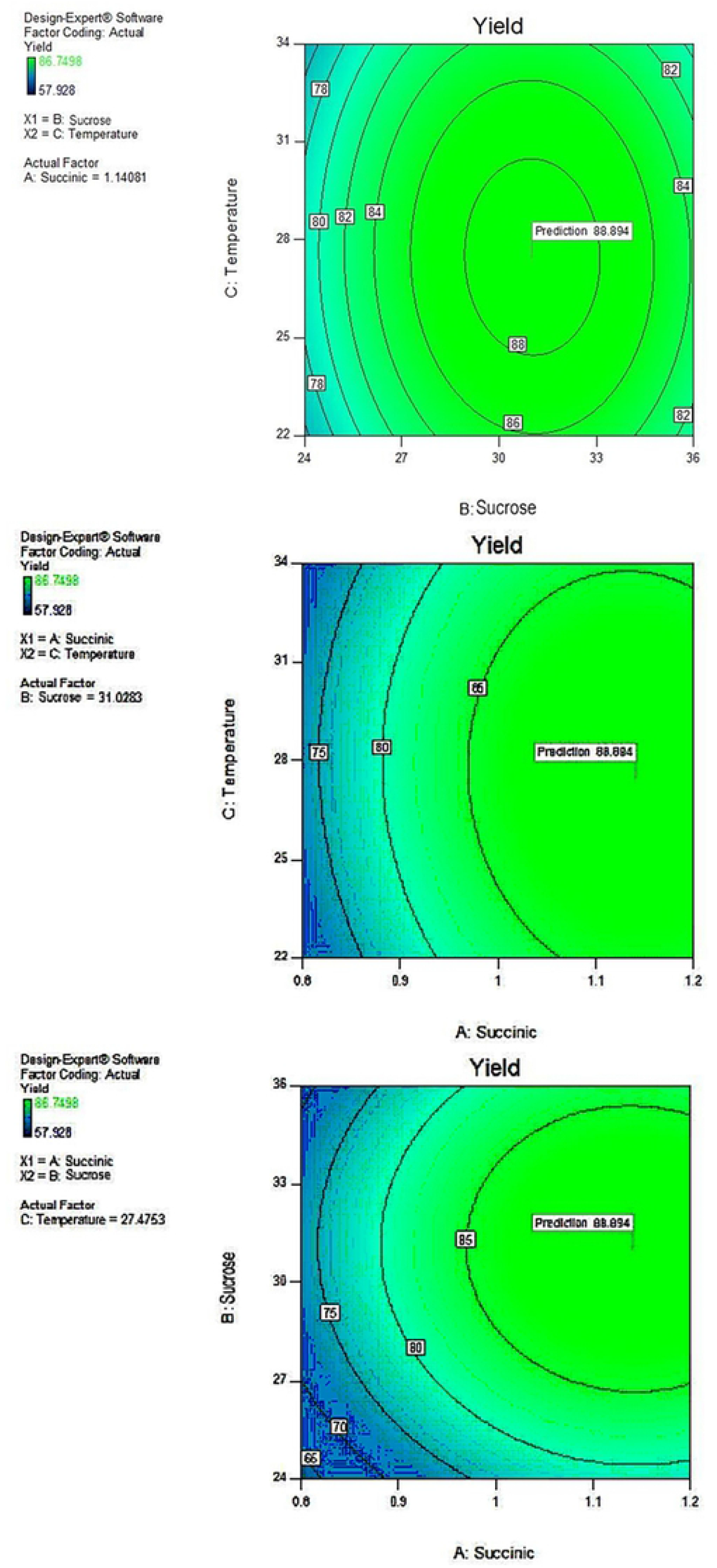
Contour plot for response surface optimization for interactions of different variables for siderophore production by *T. trachyspermus*.

**Fig 3.**
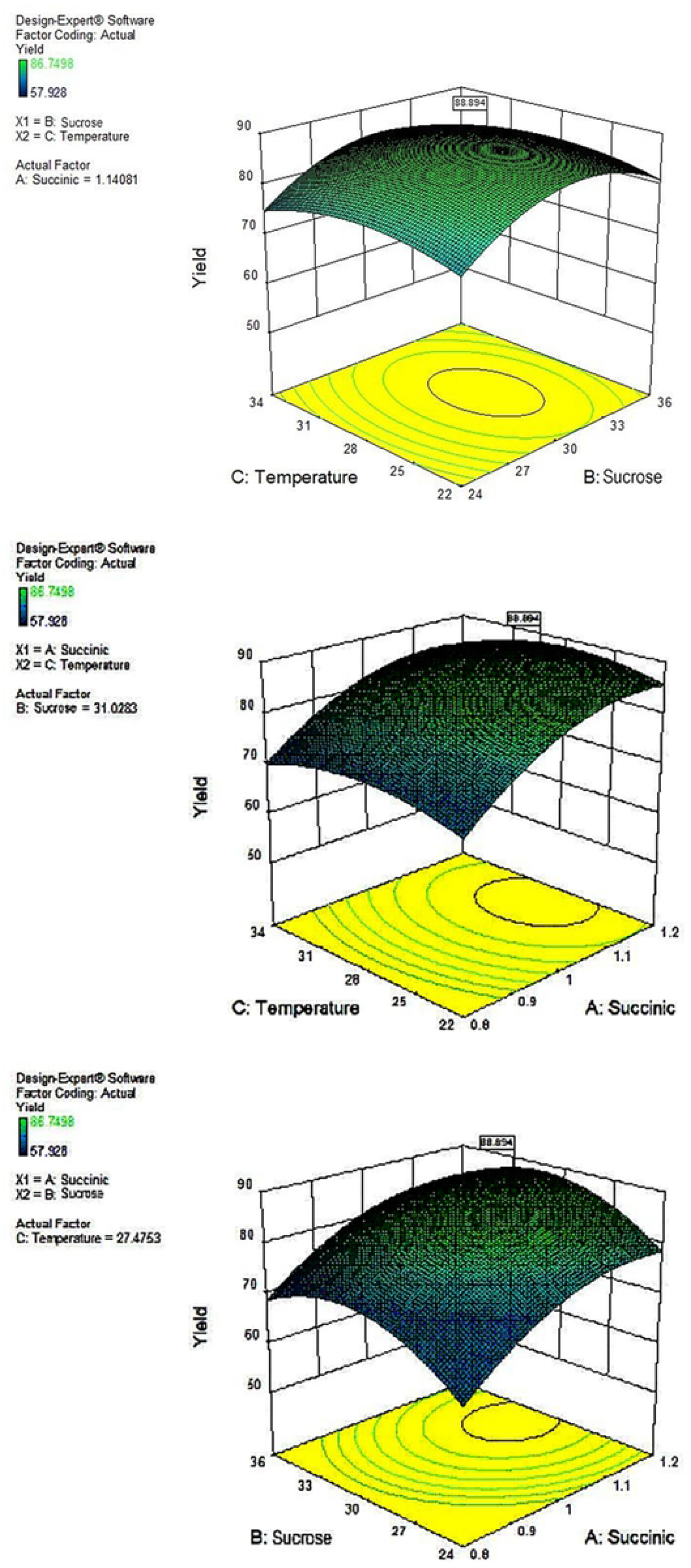
3D plot for response surface optimization for interactions of different variables for siderophore production by *T. trachyspermus*.

The maximum value of siderophore production of 88.894 obtained by maximization analysis through RSM was observed for succinic acid at a level of 1.141 g/L, sucrose concentration of 31.028 g/L and temperature of 27.475°C (Table 8) and is shown by Perturbation chart in fig 4.

**Fig 4.**
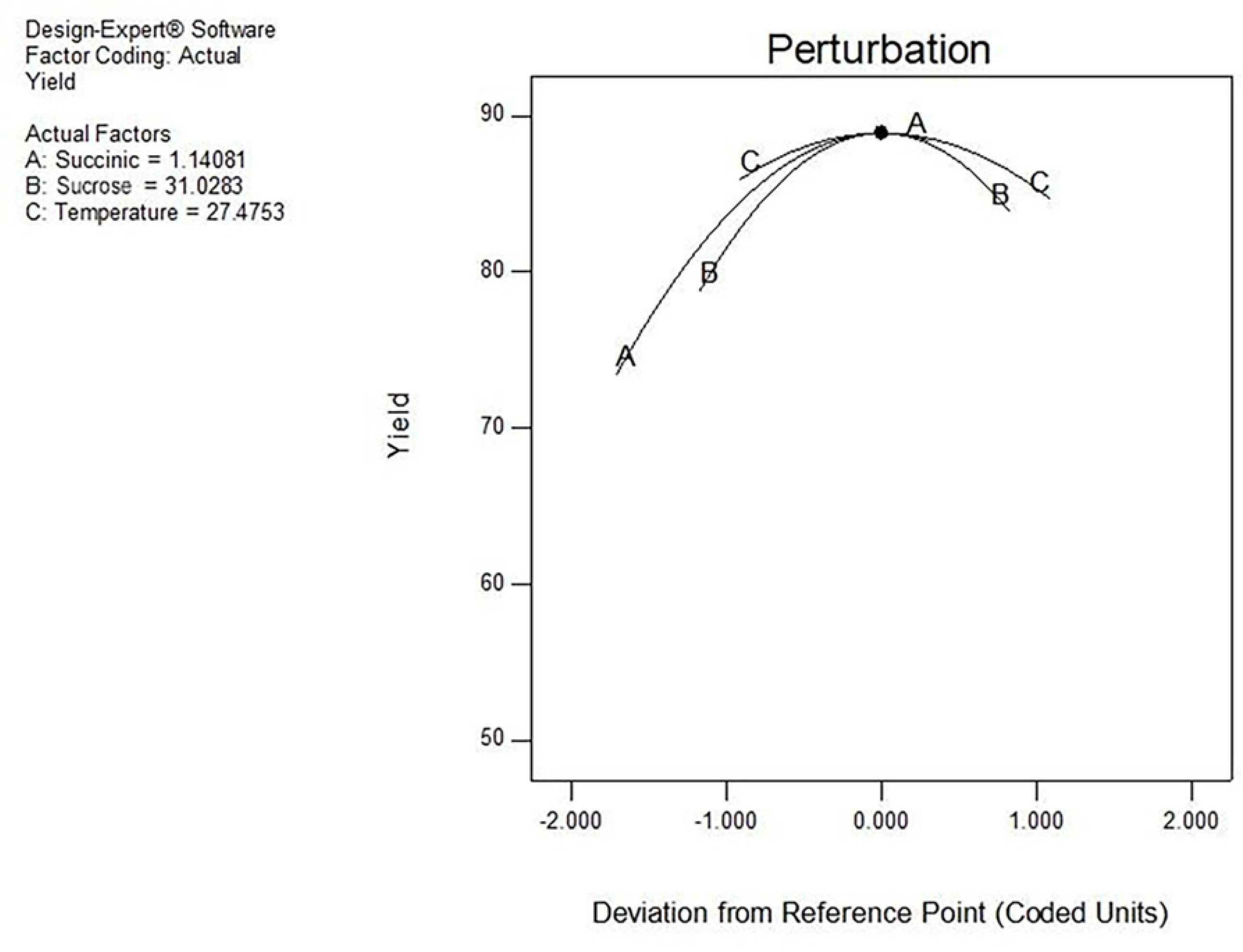
Perturbation plot for response surface optimization of siderophore production by *T. trachyspermus* as a function of succinic acid, sucrose and temperature.

The statistical optimization confers an effective and practical approach for obtaining maximum production of metabolites, Dewedar et al. [15] reported 89.33% siderophore production by *Azotobacter chrococcum* through two-steps statistical optimization method and found pH, incubation period and K_2_HPO_4_ as three significant variables. Whereas, Abo-Zaid et al. [16] reported glycerol, glucose and K_2_HPO_4_being significant variables for *Pseudomonas aeruginosa* (6.0235%) and glutamic acid, sodium succinate and FeCl_3_for *Pseudomonas fluorescens* (6.258%). 90.52% siderophore unit was achieved by using response surface methodology optimization at the glucose concentration of 21.84 g/l, pH 6.18, and Pb (NO_3_)_2_ concentration of 245.04 µmol/ by *Bacillus sp*. [17]. It was also found by Shaikh et al. that *Pseudomonas aeruginosa* produced maximum siderophore (68.41 % SU) with the optimized factors, succinic acid (0.49 g/100 ml), pH (7.08) and temperature 27.8°C [7].

In interest to our study, it is first to capitalize on the siderophore production from endophytic fungi by using the two-way statistical approach.

### Scale-up of process on bioreactor

After optimization of culture condition for siderophore production, we used fermenter (5 Liter capacity) for scale up of the production. We observed 2.98% (88.89% to 91.87% SU%) increment after 21 days of incubation in CDB medium. Results were similar to those obtained at shake flask during validation, indicating successful scale-up of process from 100 ml to 5L capacity bioreactor. Hussein and Joo have reported maximum siderophore production from *Trichoderma harzianum* (92.33%) *Aspergillus niger* (87.99%), *Metarhizium anisopliae* (85.92%) and *Penicillium digitatum* (84.26%) [18].

### Extraction and purification of siderophore

Concentrated supernatant from incubated optimized medium broth (pH 6.8) when passed through amberlite XAD-16 resin and when eluted with (1:1) methanol: water, yielded three different fractions. Out of these three fractions, only one fraction was CAS positive. This CAS positive fraction upon vacuum drying on rotatory vacuum evaporator yielded dry powder weighing 998mg/L from *T. trachyspermus* culture. This extract was further processed for characterization.

### Characterization of siderophore Detection of type of siderophores

Eluted sample of siderophore and reagent mixture were kept for 20-30 min and it was noted that pink red color in test solution appeared while no change in color was observed in control. This indicated the hydroxamate nature of siderophore compound [13] by *T. trachyspermus*. And change in the color of Arnow’s test from yellow to pink-red color indicated the presence of catecholate type of siderophore [14]. Thus, the extract contained a mixed type of siderophore as it showed presence of hydroxamate as well as catecholate type of siderophore by *T. trachyspermus*.

### Detection by Thin Layer Chromatography

When lyophilized and concentrated culture supernatant was subjected to thin layer chromatography, results were found similar to Clark and Lee [19] and Memon [20]. Thus *T. trachyspermus* produces both type of siderophores (Fig 5A and 5B).

**Fig 5.**
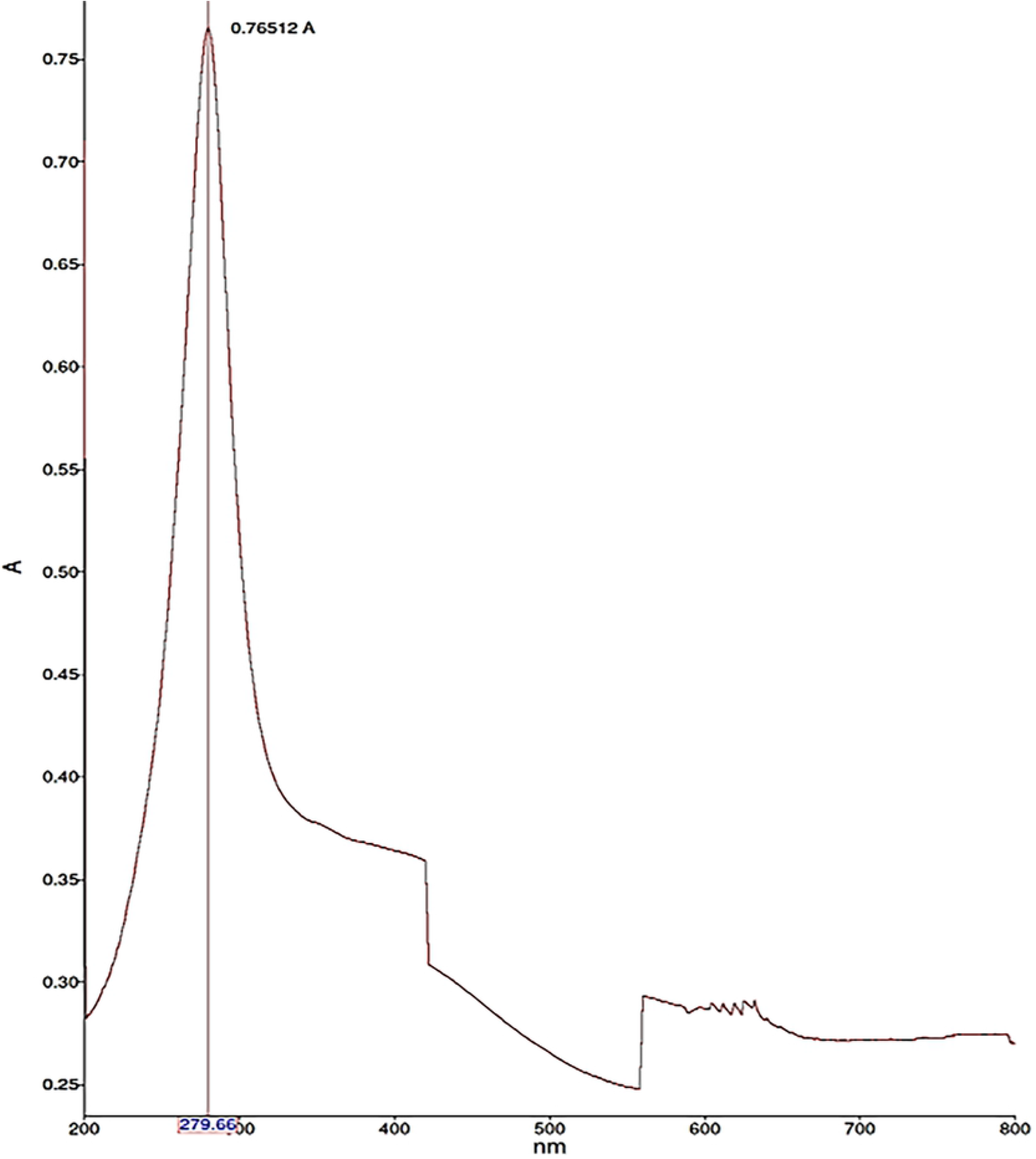
TLC plate showing both a catechol (A) and a hydroxamate-type siderophore (B) by *T. trachyspermus*.

### Spectrophotometric assay

Further, to confirm the presence of the type of siderophore in the eluted sample, spectrophotometric analysis of CAS positive fraction was carried out and found λ max at 279.66 nm (Fig 6). The compound belongs to methoxybenzene hence, the peak indicated the presence of catecholate type of siderophore [20].

**Fig 6.**
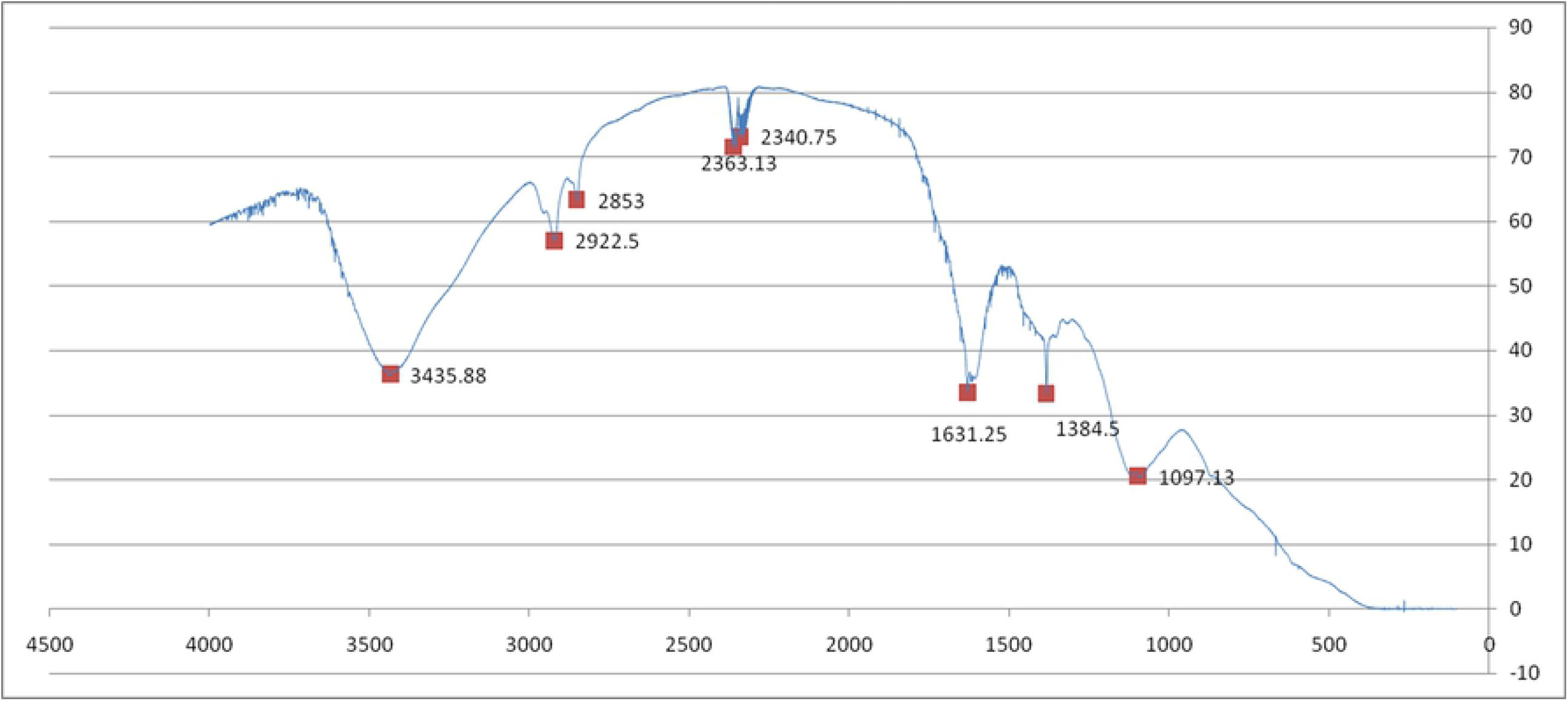
Lambda max absorption of siderophore produced by *T. trachyspermus*.

### Fourier Transform Infrared (FTIR) spectroscopy

Tank et al [21] presented that fungal hydroxamate siderophore contained functional groups like methyl, amide, secondary amine methylene, N-O bond and a ring structure (M-O) where M=Fe. Here IR spectra of mixed type of siderophore indicated the presence of hydroxyl group, carboxylic acid, amide and aromatic moieties in the molecule, appearing at 3435.88 (N-H stretch, 1°2° amines amides), 2922.5 (C-H stretch), 2853, 2363.13 (C≡N stretch, Nitriles) 2340.75, 1631.25 (N-H bend 1° amine), 1384.5 (C-H Rock, alkanes) 1097.13cm-1(C-N stretch aliphatic amines) for *T. trachyspermus* (Fig 7). This value matched with the standard values of IR frequencies. Similar results have also been reported in *P. putida* and another *Pseudomonas sp*. [8, 21].

**Fig 7.**
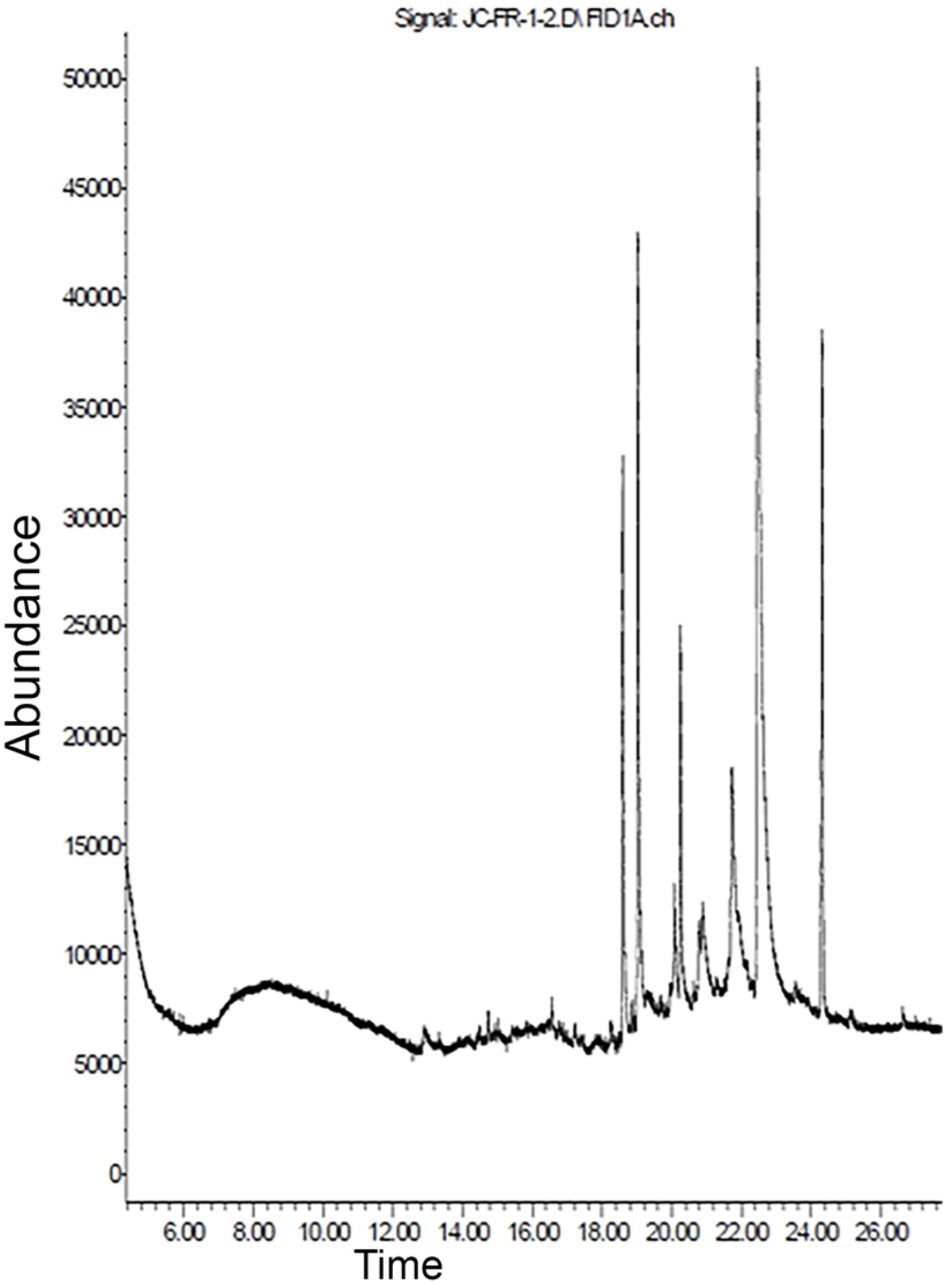
FTIR spectra for siderophore analysis produced by *T. trachyspermus*.

### GC-MS Analysis

*T. trachyspermus* produces siderophore compound having broader peak area obtainable between 22.384 to 22.859 min. of RT (Fig 8A), indicating the higher concentration of 1, 2-dihydroxybenzene (m/z-126.1) and some other compounds (M/Z-59.0, 81.0, 207.0, 281.1) (Fig 8B). Phenols with ortho-dihydroxy function can be categorized under catecholate type of siderophores [18].

**Fig 8.**
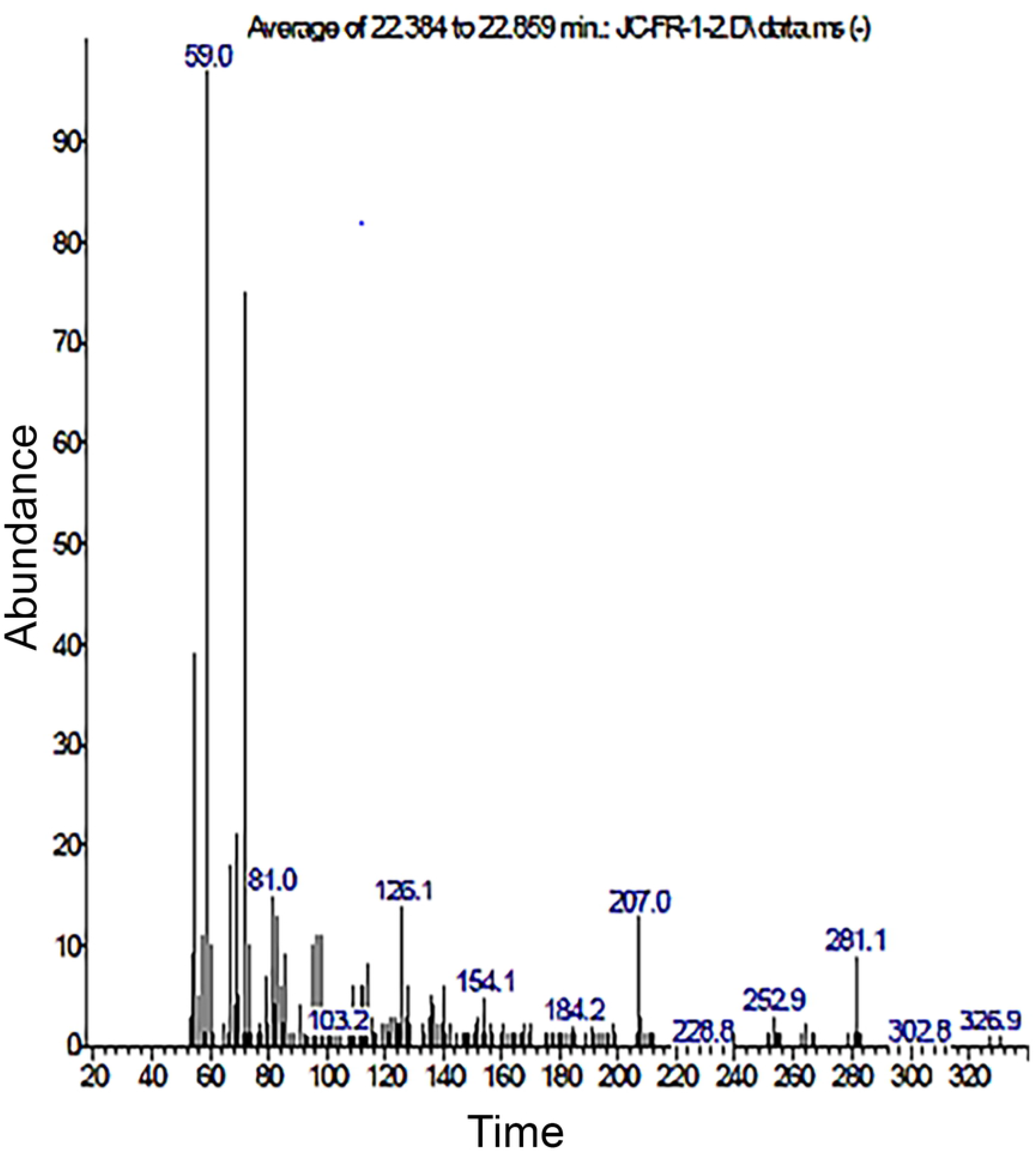
GC–MS analysis of catecholate-type of siderophores showing the GC-separation (A) and the obtained mass spectra of the separated peaks of the siderophore compound by *T. trachyspermus* (B).

### LC-MS Analysis

For determination of the identity of siderophore compound, LC-MS/MS was performed by positive electrospray ionization and with TOF acquisition SW. At physiologically relevant concentrations, by chromatogram peaks (Fig 9A) marked by blue colour at retention time 8.139 min. for berberine (9,10-Dimethoxy-2,3-(methylenedioxy)-7, 8,13, 13a-tetrahydroberbinium) with mass 335.1135 and molecular formula C_20_H_17_NO_4_ (Fig 9B), belongs to a class of alkaloid structurally related to the benzyl isoquinolines (Fig 9C), containing a second aromatic ring beside the isoquinolines as a heteropentacyclic acidic compound. This compound has been found in many plants and microbes, most notably in goldenseal (Hydrastiscanadensis), barberry (Berberisvulgaris), Oregon grape (Berberisaquifolium), and goldthread (Coptischinensis). This efficiently chelates with metal ion [22].

**Fig 9.**
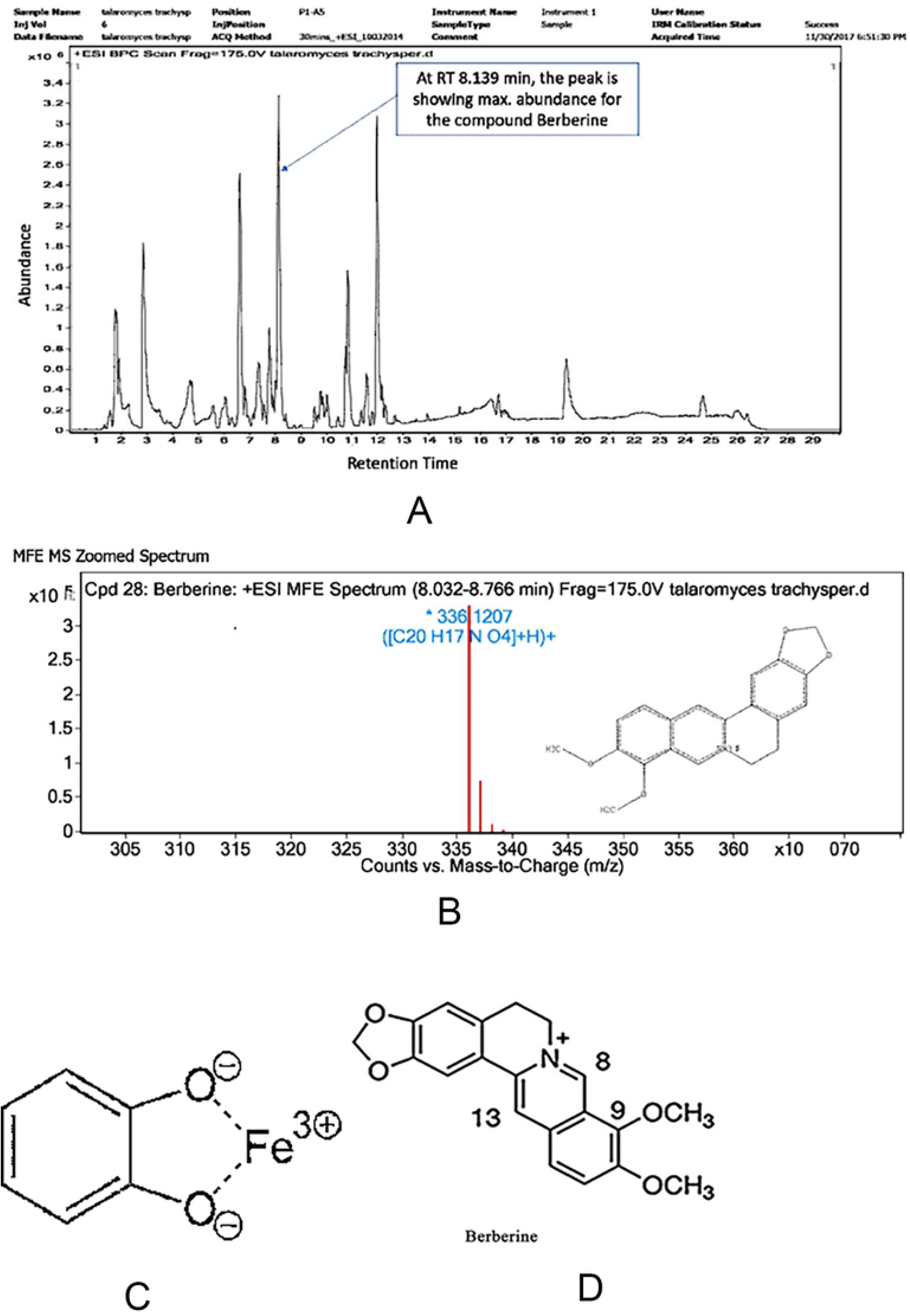
HRLC-MS chromatogram of extracted and purified siderophore compounds of *T. trachyspermus* A, MFE MS zoomed spectrum for berberine B, Catecholate-iron complex C, A detailed structure of berberine D.

Gholampour and Keikha [23] found that berberine protects the liver and kidneys against ferrous sulfate-induced toxicity by reduction in lipid peroxidation and shows ability to chelate iron. Berberine can be categorized as one of major but rare groups of siderophores, the catecholates (phenolates) [24]. Fe3+is a strong Lewis acid, preferring strong Lewis bases such as anionic or neutral oxygen atoms to coordinate with (Fig 9D) [25]. The catecholate, or catechol type, siderophores are the second most common siderophore class beside the fact that they have thus far only been found to be produced by bacteria [26]. They have been found to contain either a mono or di hydroxy benzoic acid residue that is used to chelate ferric iron and derived from di hydroxy benzoic acid [27]. The best-studied example of a catechol type siderophore is enterobactin by *E. coli* [28].

Berberine has a yellow color, and has often been used as a dye being a bioactive and very remarkable health profit phytochemical drug affecting the body at the molecular level. Berberine has been shown to reduce blood sugar [29]. Berberine has a long history of use in traditional Chinese medicine, where it was used to treat various ailments as it reduces cholesterol and triglyceride levels, while raising HDL, acts against depression, cancer, fatty liver and heart failure. It also has potent antioxidant and anti-inflammatory effects [30]. Studies have shown that it activates “metabolic master switch” an enzyme inside cells called AMP-activated protein kinase (AMPK) [31]. It has been shown to fight harmful microorganisms, including bacteria, viruses, fungi and parasites such as *Staphylococcus aureus* and *Microcystis aeruginosa* [32, 33, 34]. The endophytic fungi *Alternaria sp*., which was isolated from the healthy leaves of *Coptis chinensis* was also reported to produce 9.313 μ g• g^-1^berberine when grown in the PDA culture medium [35]. Vinodhini and Agastian reported 196 µg/L of berberine production from *Fusarium solani*, isolated from *Coscinium fenestratum* (Gaertn.) Colebr an endangered medicinal plant [36]. Galanie and Smolke reported fermentative production of protoberberine alkaloids by engineering the biosynthetic pathways in *Saccharomyces cerevisiae* [37].

## Conclusion

Today we have reached a “golden age” of metabolite reporting for the studies of several untapped biological questions in the plant and microbial sciences. Statistical optimization of physiochemical parameters of siderophore by *T. trachyspermus*, showing maximum siderophore production 88.89SU% with the prompting factor succinic acid (1.141 g/L), sucrose concentration (31.028 g/L) and temperature (27.475 °C) and on scale up production, a further increase in siderophore yield was by 3 %. (88.89 SU% to 91.87 SU %). The siderophore compound isolated from the fungi was found bioactive and characterized belonging to the rarely found catecholate group. On HRLC-MS, the compound abundantly found was named berberine, which is very imperative natural drug and has been widely reported for several medical applications. This research provided new resource for the utilization of berberine and isolation of such a fungus may provide a promising alternative approach for producing berberine.

Moreover, endophytic fungi can benefit the plant host by providing nutrients and competing with pathogenic organisms resulting in the reduction of chemical use in the agriculture, protection of agro-ecosystems and biological resources thus supporting sustainable agricultural system.

## Acknowledgement

The first author is financially supported by DBT Builder Programme Barkatullah University Bhopal (M.P.) The authors are grateful to Sophisticated Analytical Instrument Facility (SAIF) Indian Institute of Technology, Bombay for HRLC-MS/MS analysis of isolated and purified siderophore compound.

